# Quantitative genetics of lifetime growth curves in a lizard

**DOI:** 10.1101/2025.10.14.682310

**Authors:** Anaïs Aragon, Alexis Rutschmann, Murielle Richard, Elvire Bestion, Félix Pellerin, Laurane Winandy, Luis-Martin San Jose, Lucie Di Gesu, Élodie Darnet, Jehan Cribellier, Manuel Massot, Jean Clobert, Julien Cote, Pierre de Villemereuil

## Abstract

Body size is an iconic trait in quantitative genetics. For species with indeterminate growth however, growth curve, a function-valued trait, is the actual trait of interest and requires a specialised framework to study its quantitative genetics. Here, we use such a framework in the common lizard (*Zootoca vivipara*), using two large datasets (wild and mesocosm populations) with pedigrees, decomposing the total heritability in growth dynamics into two components: the heritability of “intrinsic size” (being small or large across ages) and the heritability of the shape of growth curve. We found slower growth and larger asymptotic sizes in the wild population. We show very low to moderate heritabilities in the parameters of growth curves, varying across populations and sexes. We show a small to moderate heritability in the growth curves, with a stronger importance of the heritability in the shape of growth curves than on average intrinsic sizes. Genetic variation in growth curves differed between the two populations and sexes. Our results highlight the importance of discussing the full dynamics of growth by showing that there is more adaptive potential in the earlier ages for males and in the later ages for females, with differences between the studied populations.

## Introduction

The body size of individuals impacts a large variety of their biology, from their ecology to their physiology (LaBarbera, 1989). One particular aspect of body size, for most species, is that it is generally among the most heritable traits, including in wild populations (Charmantier et al., 2014). For this reason, and also because it is generally easily measured, body size is possibly the most iconic quantitative traits (Roff, 2007). Further, due to this generally large heritable variation in body size, this trait is also expected to harbour a large adaptive potential and respond efficiently to selection, an expectation largely validated by a number of experimental evolution studies (e.g. Falconer, 1973; Partridge et al., 1999; Amaral and Johnston, 2012; Mallard et al., 2023, noting that most of these studies also confirm the tight evolutionary link between body size and life-history). While this is true for all species, species with indeterminate growth (i.e. continued growth after maturity) have an even more intricate relationship between the dynamics of their size with time and their life-history. For these species, body size is not generally considered as an actual life-history trait, yet growth is. It has a strong impact on the rest of their life-history, such as survival, age at first reproduction, competitive ability or reproductive investment (Barbault, 1988; Brown et al., 2004, 2022). This begs the following question: if body size (at any given age) tends to be heritable, what does it mean for body size integrated throughout the life-history of individuals. In other words, the question becomes: is growth itself a heritable trait? To study such question, one should stop considering “body size” as the phenotypic trait of interest, and rather consider the entirety of individual growth curves as the trait under study.

Here, we propose to study growth as a function-valued trait (Stinchcombe et al., 2012; Gomulkiewicz et al., 2018), using individual growth curves instead of instantaneous size data. Studying the quantitative genetics of such non-linear functions is not straightforward, but de Villemereuil and Chevin (2025) has recently provided a statistical and quantitative genetic framework for the study of non-linear functions in the context of phenotypic plasticity. This framework can be applied to growth curves as well, by substituting the environment for age, and estimate the full heritable genetic variation in growth strategies across individuals. Further, the method allows to disentangle heritable genetic variation stemming from *intrinsic size* (i.e. the propensity of some individuals to be larger than others, regardless of the dynamics of growth) from heritable genetic variation stemming from the *shape of growth* (i.e. variations in the shape of the growth curve itself). Such decomposition helps distinguishing static and purely morphological aspects of body size from the dynamic life-history aspects of growth throughout the lifetime of individuals. More precisely, the first component of variance (on intrinsic size) will measure heritable genetic variation between individuals in their overall tendency to be larger or smaller. By contrast, the second component (focusing on the shape of growth curves) will measure heritable genetic variation in the dynamics of individuals’ growth throughout their lifetime. Ultimately, these tools allow for a finer description of the genetic variations in growth strategies among individuals and distinguish between adaptive potential in morphology from adaptive potential in life-history through the impact of growth.

Among species with indeterminate growth, terrestrial ectotherm vertebrates such as reptiles, are of particular interest. First because growth plays a central role in their ecology (e.g., competition for territory, access to competition, Losos, 1990; Meiri, 2008). Second, because in such species growth affects size, thermoregulation and metabolism (Brown et al., 2004; Huey and Kingsolver, 2019), and therefore is a crucial part of the energetic budget that may influence the life-history strategies of individuals later in life (Metcalfe and Monaghan, 2001). Third, because ectothermic species are expected to experience an acceleration in their pace-of-life as a consequence of global warming (Bestion et al., 2015; Dupoué et al., 2022). In that regard, reptiles which specialise in cold environments may be among the most responsive, as they have evolved growth strategies that are constrained by a limited active season where individuals must trade-off energy budgets between optimizing growth, immediate survival, long-term survival and reproductive effort (Brown et al., 2004, 2022). However, little is known regarding the quantitative genetics of growth in reptiles and its variation between individuals (but see Le Galliard et al., 2006; Martins et al., 2019; Kar et al., 2023), especially in wild populations.

Here, we studied the quantitative genetics of the individual growth curves in the common lizard (*Zootoca vivipara*, Lichenstein 1823). We compared two large datasets (thousands of individuals records) of pedigreed populations: one from a cold-conditions wild population atop Mt Lozère (at 1400 m a.s.l.), and the other of a warm-conditions semi-natural mesocosm population located at the Metatron experimental facility in the South of France (400 m a.s.l.). To infer individual and genetic variation in growth curves, we used pedigree information and repeated size measurement at each age to fit a non-linear hierarchical animal model, while using the quantitative genetic framework in de Villemereuil and Chevin (2025) to relate genetic variation in the parameters of growth curves to genetic variation in size and growth.

## Methods

### Data collection

#### Studied species and populations

The common lizard (*Zootoca vivipara*) is a small ectotherm squamate specialised in cool and damp environments distributed throughout Northern Eurasia. Most populations are viviparous (Surget-Groba et al., 2006). The species is characterized by three age-classes: juveniles (zero year old), generally non-reproductive yearling (one year old) and reproductive adults (two years old or more). In viviparous populations, gestation time is of roughly two months and females lay an average of five soft-shelled eggs (one to ten), which hatch quickly after parturition.

The studied wild population is located at the top of *Mont-Lozère* (Cévennes, France), at 1400 m a.s.l., in the South-Western margin of the viviparous population distribution. Population size is fluctuating around c.a. a hundred reproducing females, with a slightly lower number of reproducing males. The number of subadults is slightly larger than the total number of reproducing adults. Each year since 1989, yearlings and adults were captured in June, soon after mating season (in May Bleu et al., 2011). The survey is still ongoing, but this study only uses data until 2019, since the pedigree information is currently lacking for the most recent years. Immediately after capture, individuals were measured, identified or marked using toe-clipping. Then, they were released at the location of capture, except for gestating females which were hosted in captivity until they gave birth (≃1 month). Within two days after birth, their juveniles were measured and marked using toe-clipping. Females and juveniles were then both released at the exact location of the female’s capture. In captivity, females were kept in individual terrariums (18 × 11 × 12 cm), filled with 3 cm of peat. A shelter was provided for thermoregulation. Terrariums were placed with one corner below a 25 W heating bulb (active 6 hours a day: 9 am – 12 pm / 2 pm – 5 pm) to create a thermal gradient. The room followed an artificial 10 h–14 h day-night cycle starting from 2003 and natural day-night cycle before that (roughly 15 h–9 h). Terrariums were humidified three times a day and females were fed with two medium-size house crickets (*Acheta domesticus*) every two days.

The semi-natural mesocosm lizard population was made of lizards inhabiting 16 mesocoms in the Metatron facility, at the Station of Theoretical and Experimental Ecology (Ariège, France, 43°01’ N, 1°05’ E).The Metatron is an enclosure system composed of 48 mesocosms of 100 m^2^ each hosting a diverse community of plant and invertebrate species similar to natural wetland habitats (Legrand et al., 2012; Bestion et al., 2015). Each mesocosm is fully enclosed by underground tarpaulins and fine-meshed nets to prevent species to enter and exit. The lizard population was established in 2010 for the purpose of warming experiments (Bestion et al., 2015; Pellerin et al., 2022). The lizards used in the experiments were descendants of lizards captured in the Cévennes (i.e. in the same region as the wild population) in 2010, 2012 and 2014. There are on average c.a. 140 reproducing females in the populations, with the same proportion of males and sub-adults as described above for the wild population. The monitoring and hosting procedures were similar to the ones used in the natural population and were performed annually from 2010 to 2022.

Despite the similar regional origins of both populations, given the location difference between the populations, the environment was clearly different between both populations. This includes multiple factors, such as climate or plants and insects communties. Differences in the thermal environment are notably striking: the mesocosm at Metatron was a warmer environment, with an average summer temperature of 19.0 °C, while the wild population had an average summer temperature of 13.7 °C.

#### Phenotypic data

Snout-vent length (SVL), i.e. the length from the extremity of the snout to the cloaca (Garland, 1988; Sorci and Clobert, 1999; Sion et al., 2021, a robust and common measure of body size in squamates), was measured for all individuals (juveniles and individuals caught in the field) using a transparent ruler and gently stretching individual alongside it. The reproductive stage (juvenile, non-reproductive yearling, reproductive adult of age ≥ 2), as well as the sex of all individuals, were recorded. For the juveniles, the sex was inferred from counting ventral scales, as individuals with more ventral scales are more likely to be females (Lecomte et al., 1992). Individuals were subsequently sexed as yearlings and/or adults, and if necessary sex was corrected, as errors became unlikely as individuals grew older due to sexual dimorphism. The sample sizes for the different models used in this article are provided in Table 1.

**Table 1.** Sample size table for both populations, when performing model selection (Full) or the quantitative genetics analysis (Animal model). *N* is the total sample size and *N* Ind. is the number of individuals. Sample sizes vary due to juvenile mortality (e.g. *L*_0_ v. *k* and *L*_*∞*_) or pedigree information availability (e.g. full model v. animal models in the wild population).

|  | Wild |  | Enclosed |  |
| --- | --- | --- | --- | --- |
| | $N$ | $N$ Ind. | $N$ | $N$ Ind. |
| **Full (model selection)** | 6714 | 2809 | 2734 | 972 |
| **Animal model ($L_0$)** | 5404 | 5404 | 4033 | 4033 |
| **Animal model ($k$ and $L_\infty$)** | 3176 | 1231 | 2734 | 972 |

#### Pedigree reconstruction

For each new individual (caught in the field or born in the lab), a tail tip tissue (*<* 1 cm for yearlings or adults, *<* 3 mm for juveniles) was collected (tail grows back in this species). DNA was extracted from tail tissue using DNeasy Blood & Tissue (Qiagen) purification kits. We then genotyped all individuals at 8 microsatellite loci (2 different sets for the wild or mesocosm population). The CERVUS software (Marshall et al., 1998) was used to reconstruct paternities of juveniles born in the lab (20% of missing parternities for both populations). Most maternities were known from observation in the breeding facilities. They could also be reconstructed for juveniles born in the field for the mesocosm population. The maximum pedigree depth was 8 in both populations, with mean depth of 2.24 and 2.79 for the wild and mesocosm populations respectively.

#### Ethics statement

Captures were allowed by the French DREAL (Licence no. 2010-189-16, 2013-274-0002 & 2021-s-22) and measurements were performed under the approval of the Cuvier Ethics Committee (authorisation number APAFIS#15897-2018070615164391, APAFIS#19523-201902281559649 v3 & APAFIS #34093-2021112211094302 v4) in animal breeding facilities accredited with permit DDETSPP-SPAE-2022-262-001 for the wild population and B09583 for the mesocosm population.

### Statistical analysis

#### From size series to growth curve

To analyse these series as growth curves, i.e. function *f* (*t*) yielding length *L*_*t*_ at age *t*, we first investigated the fit between our data and three different choices of growth curves classically used in the literature (Schoener and Schoener, 1978; Johnston, 2011; Hernández-Salinas et al., 2019; Yang et al., 2019; Rotger et al., 2023): *(i)* a von Bertalanffy growth curve:

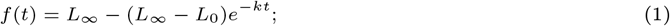

*(ii)* a logistic-by-length growth curve:

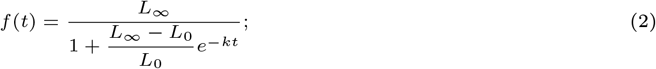

and *(iii)* a Gompertz curve:

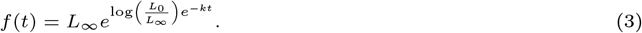

In each of these models, *L*_0_ is the SVL at birth (*t* = 0), *L*_*∞*_ is the asymptotic SVL at old ages and *k* is a growth coefficent determining the “speed” of growth (it is the only parameter related to *t*), notably the overall speed at which the asymptotic plateau is reached. The parameters are described in Table 2. The impact of variation of the three parameters on the overall shape of the curves is illustrated in Figure 1. The differences between the three growth models are detailed in Schoener and Schoener (1978) and Yang et al. (2019). Briefly, while the logistic growth curve assumes a linear decrease of relative growth with size, van Bertalanffy assumes an accelerated decrease, and Gompertz a negative logarithmic decrease of growth with size. Note also that in all models, *L*_*∞*_ is an asymptotic size (individuals are assumed to technically keep growing, only less and less), which does not correspond to size at maturity (reached around age 2, frequently age 1 in the mesocosm population), arising from allometric and metabolic constraints on growth (Schoener and Schoener, 1978). The fitting and model selection are described below. For the mesocosm data, given the considerable spread in dates of birth, we used a measure of age precise to the day. This was not possible for the wild population because many date of birth are missing for animals captured as yearlings or adults for the first time. Yet the spread of birth dates is less important there which minimises this problem (s.d. of 3.9 v. 11.1 for the mesocosm population).

**Table 2.** Lexical table of terms and parameters used in the article.

| Name | Notation | Explanation |
| --- | --- | --- |
| Size at birth | $L_0$ | Growth curve parameter: size of individuals at birth. |
| Asymptotic size | $L_\infty$ | Growth curve parameter: asymptotic size defining the plateau of size reached at old ages. |
| Growth coefficient | $k$ | Growth curve parameter: coefficient relating time (years) with the speed of growth, defining the overall speed at which the asymptotic plateau is reached. |
|  | • | Stands for any of the growth curve parameter above. |
| Additive genetic variance of parameter | $V_{A,\bullet}$ | Additive genetic variance of parameter • inferred from the animal model. |
| Maternal variance of parameter | $V_{M,\bullet}$ | Maternal effect variance of parameter • inferred from the animal model. |
| Year variance of parameter | $V_{Y,\bullet}$ | Yearly effect variance of parameter • inferred from the animal model. |
| Fixed-effect variance of parameter | $V_{F,\bullet}$ | Variance arising from the fixed-effects explaining parameter •, as inferred from the animal model (see de Villemereuil et al., 2018). |
| Permanent environment variance of parameter | $V_{PE,\bullet}$ | Permanent environment effect variance (see Wilson et al., 2010) of parameter • inferred from the non-genetic individual effect in the animal model (only applicable to $k$ and $L_\infty$). |
| Residual variance of parameter | $V_{R,\bullet}$ | Residual variance of parameter • in the animal model (only applicable to $L_0$). |
| Total variance of parameter | $V_{Par,\bullet}$ | Total variance in parameter • inferred from the animal model. |
| Heritability of parameter | $h_\bullet^2$ | Heritability of •, i.e. $V_{A,\bullet}$ divided by $V_{Par,\bullet}$. |
| Total variance among individuals in growth | $V_{Tot}$ | Total variance of individuals in their growth dynamics across all ages. |
| Total additive genetic variance in growth | $V_{Add}$ | Total heritable genetic variance contained in the individual growth curves across all ages. |
| Heritability of growth | $h_{Growth}^2$ | Total heritability in the growth dynamics, i.e. $V_{Add}$ divided by $V_{Tot}$. |
| Additive genetic variance in intrinsic size | $V_A$ | Heritable genetic variance due to variation in average size (across ages) among individuals, i.e. their heritable variation in being larger or small than others consistently across ages. |
| Heritability of intrinsic size | $h_{Size}^2$ | Heritability in the intrinsic size, i.e. $V_A$ divided by $V_{Tot}$. |
| Additive genetic variance in the shape of growth | $V_{AxAge}$ | Heritable genetic variance due to individual differences in the shape of the growth curves, i.e. their heritable variation in the dynamics of growth (rather than in intrinsic size). |
| Heritability of growth shape | $h_{Shape}^2$ | Heritability in the shape of the growth curve (in growth dynamics), i.e. $V_{AxAge}$ divided by $V_{Tot}$. |
| γ-decomposition | $\gamma_\bullet$ | Decomposes the contribution of the heritable variation of each parameter • to the total heritable variance $V_{Add}$. Sums up to 1 and be performed across all ages, or at any given age. |
| ι-decomposition | $\iota_\bullet$ | Decomposes the contribution of the heritable variation of each parameter • to the heritable variance in growth shape $V_{AxAge}$. Sums up to 1 and can only be performed across all ages. |

**Fig. 1.**
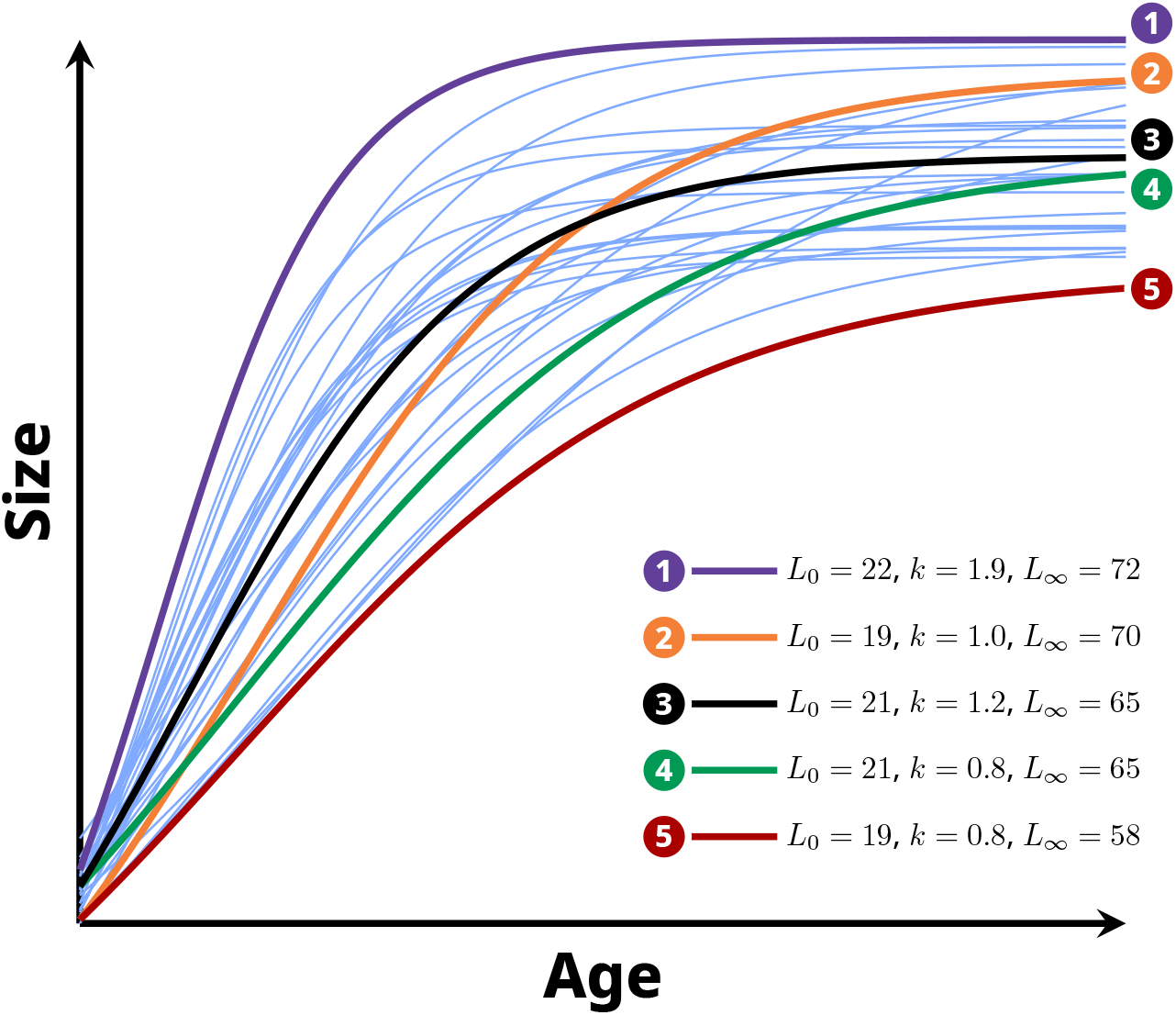
Fictional growth curves with parameters close to the values observed on the common lizard. The blue thin lines are individual growth curves. Other curves (purple 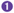, orange 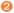, black 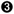, green 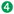, red 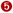) illustrate particular cases with their parameters displayed in the legend. The line 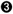 also happens to be an average growth curve. Parameters are size at birth *L*_0_, growth coefficient *k* and asymptotic size *L*_*∞*_. The 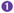 and 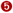 lines show individuals with large (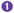, always above the others) or small (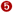, always below the others) intrinsic sizes, i.e. average sizes across life. The 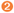 curve shows an individual with a strong difference in the shape of growth compared to the 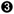 curve, while having a roughly average intrinsic size (roughly same average size as 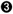). The 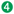 curve demonstrates the influence of parameters across ages: it has the same growth coefficient as 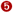, showing that size at birth influence size at later stages and that curves with equal growth coefficients are “parallel” only to a limited extend, 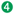 and 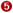 curves have a slightly different shapes despite having the exact same growth coefficient. These particular cases show the difficulties in interpreting differences in individual growth trajectories directly from parameter values differences.

#### Imputation of missing birth size

For the wild population, when individuals were caught as juvenile or yearling for the first time, their age was known and we only missed their size at birth, which corresponds to the parameter (*L*_0_) of our model. Given the low variability of size at birth (coefficient of variation of 0.057), and the relatively high number of individuals caught as yearlings, we imputed their size at birth, to be able to analyse the rest of their SVL series. We verified that removing individuals with imputed size at birth did not impact model selection. To do so, we used multiple imputation (Rubin, 1987) and the mice R package (van Buuren and Groothuis-Oudshoorn, 2011) to produce 20 imputed datasets, providing sex, year of birth, size at first year and month of capture at first year as codependent variables for imputation. Individuals caught as adults for the first time were excluded due to missing (and not imputable) age. For the mesocosm dataset, the yearly effort of capture being much more important (10 sessions a year) and the probability of capturing all individuals being higher in the mesocosm population, such imputation was not necessary.

#### Model fitting and growth curve selection

Using a Bayesian framework and brms R package (Bürkner, 2018), we fitted the models in Equation 1, Equation 2 and Equation 3 as non-linear models of size against age. Models were compared using their widely applicable information criterion (WAIC) (Vehtari et al., 2017). We selected individuals for which at least two records were available (we also checked with three records, with no improvement on the credible intervals of the estimates), resulting in 6714 measures on 2809 individuals for the wild population and 2734 measures on 972 individuals for the mesocosm population. Because WAIC is not defined for multiple imputed models, we provided the average imputed values for the model selection phase for the wild population, then the best model was re-fitted using multiple imputation. We defined informative, but wide, priors based on previous studies providing information on the size-age relationship of the common lizard (Heulin, 1985; Le Galliard et al., 2006; Horváthová et al., 2013a; Clobert et al., 2014). For each model, 10 chains were run for 3000 iterations, with a warm-up of 1000 iterations. The final, best model was run with a unique chain of 2000 iterations with a warm-up of 1000 iterations and thinning of 5. Effective sample size (ESS > 800) and 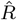 *<* 1.05 (Vehtari et al., 2021), as well as graphical inspection of the traces of parameters, were used to diagnose issues with the algorithm, except for the final model using multiple imputed datasets, which impacts the validity of these statistics.

#### Quantitative genetics model

We used the best-fitting curve *f* (*t*) for growth to fit a non-linear hierarchical animal model, i.e. a model where the non-linear curve parameters *k* and *L*_*∞*_ were allowed to vary across individuals, according to their own identity, year of birth, maternal identity and sex. Individual identity effects were split between an additive genetic effect and a purely independent individual-level effect (“permanent environment” effect, Wilson et al., 2010):

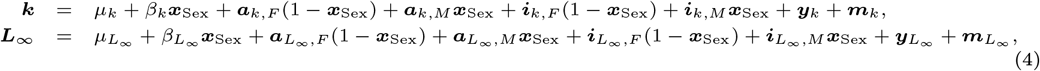

where *μ*_*•*_ is the population-level (female-specific) value of the parameter for parameter • (*k* or *L*_*∞*_), *β*_*•*_ is the male-specific fixed effect (with ***x***_Sex_ being 0 for female and 1 for male), ***a***_*•*,*S*_ is the additive genetic random effect for sex *S* (*F*: female, *M*: male), ***i***_*•*,*S*_ is the individual-level random effect for sex *S*, i.e.`’permanent environment” effect, ***y***_*•*_ the year-of-birth random effect and ***m***_*•*_ is the maternal random effect. These variances are described in Table 2. Additive genetic effects were modelled as following a multivariate normal distribution depending on the relatedness matrix, ***A***, obtained from the pedigree (Henderson, 1950; Kruuk, 2004):

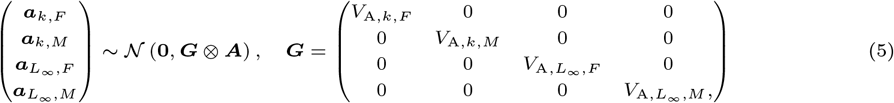

where ⊗ is the Kroenecker product and ***G*** is the additive genetic variance-covariance matrix, with *V*_A,*•*_ being the additive genetic variance of parameter •. We did not infer the additive genetic covariance between *k* and *L*_*∞*_, because preliminary models showed we lacked power to do so and it considerably slowed down the algorithm (note that this assumption does not influence the variance inference). All the other random effects were modelled as independent, following a normal distribution of variance respectively *V*_PE_, *V*_Y_ and *V*_M_ for *i*_*•*_, *y*_*•*_ and *m*_*•*_.

Since the size at birth is both a parameter of the model (*L*_0_) and an observed value (even part of the response variable), it was impossible to include effects on its in the hierarchical part of the model, due to limitations in the syntax of brms. Instead, size at birth was considered as a directly observed parameter *L*_0_ in the non-linear model, and we analysed its variation in a separate linear animal model. As we did not fit the covariance between the different parameters, this two-step approach did not impact our ability to combine all estimations in a common **G**-matrix for the three parameters. In doing so, we neglect the inference of the additive genetic covariance between parameters, which was already absent of the inference for *k* and *L*_*∞*_ and does not impact the inference of the variances themselves. Separated inference between sexes for the additive genetic variance in the model for *L*_0_ did not yield different additive genetic variances between sexes and thus a unique additive genetic variance was inferred for this parameter. Apart from this absence of sex differences, the same model was used for *L*_0_ as for *k* and *L*_*∞*_ in Equation 4. We added the date of birth (number of days since the first of June) as a fixed effect for *L*_0_.

Models were run using the brms package (Bürkner, 2018). The non-linear animal model, for *k* and *L*_*∞*_, was run for 2000 iterations, with a warm-up of 1000 iterations, one chain for each of 20 imputed datasets. The linear model for *L*_0_ was run with 10 chains of 3000 iterations, with a warm-up of 1000 iterations. Since posterior medians are the best point estimates for variances (Pick et al., 2023) and generally not very different from the mean for fixed effects, we report it for all estimates, along with the 95% credible intervals (CI) computed from the posterior quantiles. All *p*-values provided are Bayesian *p*-values, which have asymptotic properties akin to frequentist *p*-values (Shi and Yin, 2021).

#### Variance decomposition

To compute and decompose the additive genetic variance contained in the growth curve, we applied the framework described by de Villemereuil and Chevin (2025) and the Reacnorm package. While developed in the context of reaction norms, the genetic decomposition part of the framework for de Villemereuil and Chevin (2025) is a general way to compute and decompose the additive genetic variance of any non-linear function-valued trait, by relating heritable variation in the parameters of the function to the scale of the trait. Such relation is based on the additive properties of heritable genetic variation, using the linear average gradient of the function according to all parameters (see de Villemereuil and Chevin, 2025, for more details). The Reacnorm package allowed us *(i)* to compute the additive genetic variance *V*_Add_ of “growth” (i.e. contained in the growth curve); *(ii)* to separate *V*_Add_ into the additive genetic variance in “intrinsic size” (i.e. due to genetic differences between individuals in their average size along their life, *V*_A_) and the additive genetic variance in the shape of growth curve (i.e. due to additive-by-age interaction, *V*_AxAge_); and *(iii)* to decompose *V*_Add_ (*γ*-decomposition) and *V*_AxAge_ (*ι*-decomposition) into the relative contribution of each of the three parameters of the model (*L*_0_, *k, L*_*∞*_). From the additive genetic variances and the total variance (*V*_Tot_, excluding the variance arising from the average shape of the growth curve, i.e. conditional to the average impact of age), we computed the heritability of the growth curve 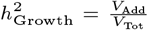, the heritability of intrinsic size (i.e. of being on average larger or smaller than other individuals across ages) 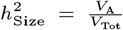 and the heritability in the shape of the growth curve 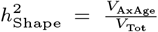. Since *V*_Tot_ is a common denominator, this follows the decomposition of the additive genetic variance described above and results in the following decomposition of heritability of growth:

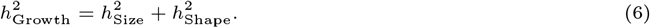

These variances and heritabilities are described in Table 2.

To understand what we mean by “intrinsic size” and its heritability, compare the extreme (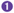 and 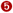) growth curves in Figure 1: the individual with the growth curve 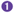 always has a larger size than other individuals, while the 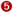 one always has a smaller size. The intrinsic size of individual 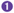 is thus the largest among individuals, while it is lowest for 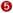. Any heritable variation in the set of parameters resulting in such kind of consistent variation in size across ages will contribute to increasing the value of 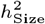. On the contrary, comparing the growth curve 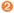 to 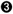 in Figure 1 show that, despite having smaller size at earlier ages, 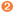 individual tend to have a larger size at later ages, with a roughly equal intrinsic size compared to an individual following the growth curve 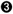. In this case, the distinction between 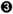 and 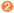 individuals is not their intrinsic size, but rather the dynamics of their growth. In other words, their differ in the “shape” of their growth curve. Any heritable variation in the parameter resulting in such kind of variation in shape will contribute to increasing the value of 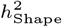.

The *γ*- and *ι*-decompositions offer to go deeper into the relationship between genetic variation in the growth curve parameters (*L*_0_, *k, L*_*∞*_) and the function-valued trait additive genetic variances (*V*_Add_ for *γ*-decomposition and *V*_AxAge_ for the *ι*-decomposition). In other words, a large value for a parameter in the *γ*-decomposition mean that genetic variation in this parameter contribute mostly to the total heritable variance. The *γ*-decomposition can also be performed age-by-age, showing the relative contribution of the genetic variation in parameters at a given age (see Figure 1 for explanation of how parameters influence different stages of life). When *γ*-decomposition value of a parameter is high at a given age, it means that response to selection at this particular age will mostly rely on genetic variation in this parameter. Age-by-age computation was only performed from age 0 to 6, because after ages above 6, the heritabilities and *γ*-decomposition became stable, reflecting the asymptotic plateau reached by the model. Because it targets heritable variation in the overall shape of growth curve (*V*_AxAge_ and the related heritability 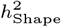), the *ι*-decomposition cannot be performed age-by-age, but must be performed overall. As for the *γ*-decomposition, a high value for a parameter in the *ι*-decomposition means that heritable genetic variance in this parameter contributes most to most of the heritable genetic variation in the shape of growth curve (i.e. to *V*_AxAge_). This means that response to selection on the *shape* of growth curve will mostly rely on heritable variation in this parameter.

## Results

### Growth curve selection

For both the wild and mesocosm populations, the model with the best fit was the logistic growth model, with a large difference in WAIC compared to the second best model (Table 3). The logistic growth curves for males and females, in the wild and mesocosm populations, are shown in Figure 2. For all ages, males were on average smaller than females. In the wild population, the asymptotic plateau was reached near the fourth year, with an asymptotic size near 65mm for the females and close to 60mm for the males. By comparison, individuals from the mesocosm population reached their asymptotic plateau sooner, near their second year. Their asymptotic size was also slightly smaller, with slighlty stronger sexual dimorphism (near 63mm for females v. 56mm for males).

**Table 3.** Table providing the Widely Applicable Information Criterion (WAIC) for the two datasets (wild and mesocosm populations) and the three shapes of growth curve tested: a von Bertalanffy curve, a logistic-by-length curve and a Gompertz curve. The best fitting curve has the lowest WAIC.

| Model | Wild |  |  | Mesocosm |  |  |
| --- | --- | --- | --- | --- | --- | --- |
| | WAIC | ΔWAIC | Std. Error | WAIC | ΔWAIC | Std. Error |
| **Logistic** | 32542.3 | 0.0 | 0.0 | 8968.2 | 0.0 | 0.0 |
| **Gompertz** | 33933.7 | -695.7 | 28.2 | 9325.6 | -178.7 | 25.8 |
| **Von Bertalanffy** | 35273.4 | -1365.6 | 50.3 | 10074.6 | -553.2 | 51.0 |

**Fig. 2.**
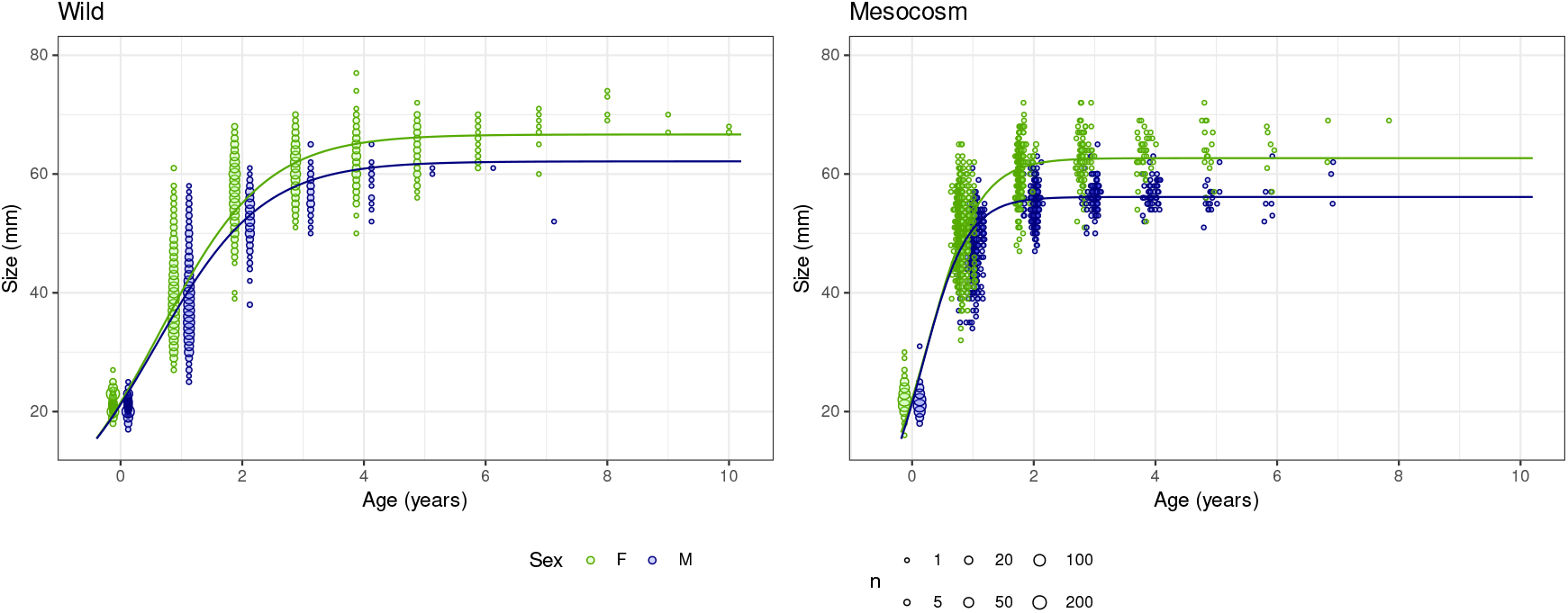
Growth curves (size against age) for males (blue) and females (green) for the logistic model, for the wild and mesocosm populations. Actual data are shown in the background for males (pale blue) and females (pale green), with dot size reflecting the number of individuals sharing the same couple of size and age. Location of the points reflect the minimal precision of age (to year for the wild population and to the day for the mesocosm population) and are slightly shifted between males and females for better readability.

### Size at birth

The size at birth (*L*_0_) was comparable between the wild and mesocosm populations, between 21 and 22 mm (Table 4). It differed between sexes, with males being shorter than females on average (wild: Δ = −0.637 mm [−0.681 mm, −0.592 mm], *p <* 5e-5; mesocosm: Δ = −0.684 [−0.741, −0.626], *p <* 5.00e − 5). It also depended on the birth date (wild:*b* = −0.0171 [−0.0264, −0.00799], *p* = 1.00e-4; mesocosm: *b* = −0.0267 [−0.0305, −0.0228], *p <* 5.00e-5), with earlier juveniles showing larger size. For both populations, the most important explanatory factor of size at birth was the cohort effect (*V*_Y_, Table 4), reflecting the strong impact of the current environmental conditions on the ability of females to invest in their offspring size. This effect was followed by maternal, then additive genetic, effects (*V*_M_ and *V*_A_, Table 4). However, both *V*_M_ and *V*_A_ contributed more to explain the total variance in the mesocosm population, compared to the wild population. Consequently, heritability of size at birth was small, but with support away from zero, in the wild population 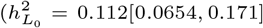, Table 4), and substantially larger in the mesocosm population 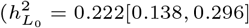, Table 4).

**Table 4.**
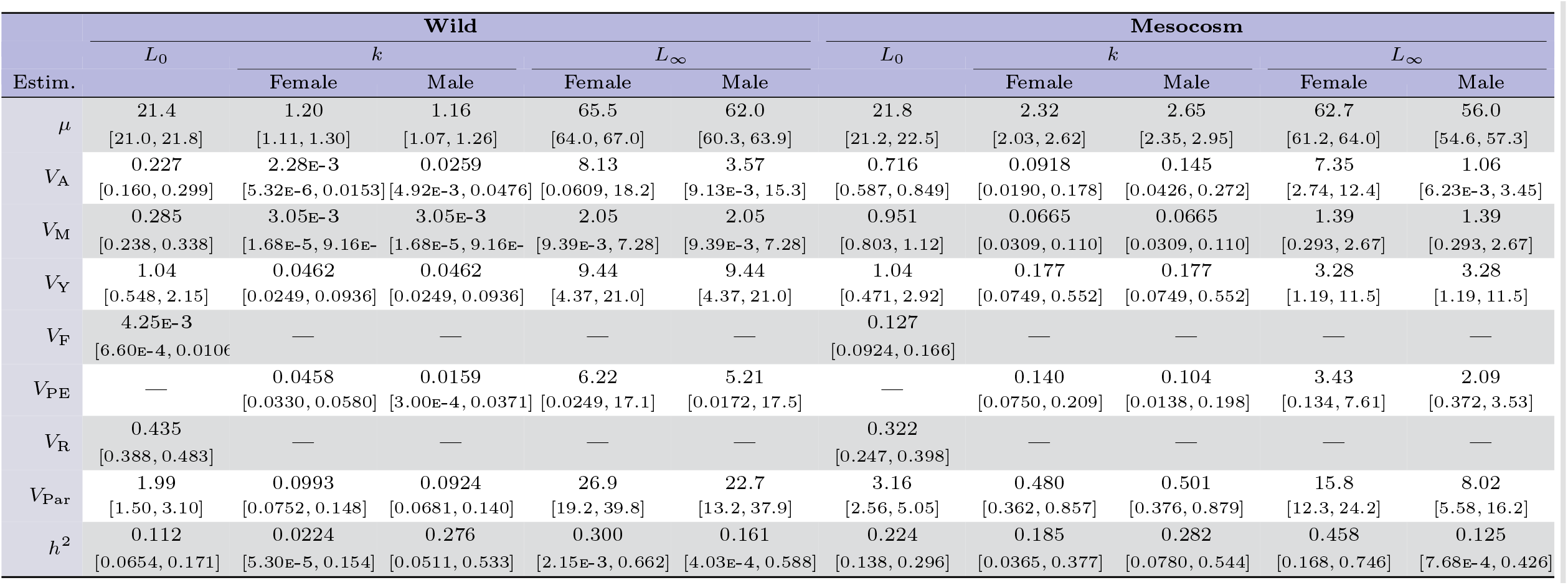
Estimates of the phenotypic mean (*μ*), additive genetic variance *V*_A_, maternal effect variance (*V*_M_), cohort effect variance (*V*_Y_), fixed-effects variance (*V*_F_, without the effect of sex), permanent environment effect (*V*_PE_) and residual variance (*V*_R_) for each of the three parameters of the model, from both the wild and mesocosm populations. For *L*_0_, estimates come from the linear animal model with size at birth as a response variable. For *k* and *L*_*∞*_, estimates come from the non-linear animal model with size at different ages as a response variable. The total variance in each parameter (*V*_Par_) is the sum of all the variance components for that parameter. For non-linear model parameters, the (intra-individual) residual variance is replaced by a permanent environment effect which plays virtually the same role. All estimates are provided here conditional to the effect of sex.

### Parameters of growth curve

The growth coefficient *k* was twice as small in the wild population compared to the mesocosm population (Table 4). In both cases, there was considerable variation among individuals depending mainly on their identity and year of birth (*V*_PE_ + *V*_A_ and *V*_Y_, Table 4). Maternal effects were still a substantial factor (*V*_M_, Table 4), especially in the mesocosm population. There was no difference in the average growth coefficient between males and females in the wild population (Δ = 0.0389 [−0.0843, 0.000670], *p* = 0.0935), but the growth coefficient was larger for males in the mesocosm population (Δ = 0.328 [0.221, 0.436], *p <* 5.00e-5). By contrast, the additive genetic variance was ten times larger for males than for females in the wild population (log10-ratio = 1.03 [6.15e-4, 3.62], *p* = 0.0498), but not in the mesocosm population (log10-ratio = 0.200 [−0.354, 0.865], *p* = 0.408). The resulting heritabilities followed the same pattern, with heritability being ten times larger for males than for females in the wild population (log10-ratio = 1.06 [0.0484, 3.65], *p* = 0.0396). Male heritability was also higher in the mesocosm population, but not significantly so (log10-ratio = 0.184 [−0.347, 0.829], *p* = 0.416).

The average asymptotic size *L*_*∞*_ was larger for the wild population compared to the mesocosm population (Table 4). Males had significantly smaller asymptotic size than females for both population (wild: Δ = −3.50 mm [−4.88 mm, −2.01 mm], *p <* 5.00e-5; mesocosm: Δ = −6.67 mm [−7.35 mm, −5.99 mm], *p <* 5.00e-5). There was substantial variation among individuals, again depending mainly on their identity and year of birth (*V*_PE_ + *V*_A_ and *V*_Y_, Table 4). Maternal effects were small compared to other effects acting on the asymptotic size variation (*V*_M_, Table 4). In both populations, the additive genetic variance was smaller in males compared to females, although the discrepancy is significant only in the mesocosm population (wild: log10-ratio = −0.300 [−2.88, 1.75], *p* = 0.612; mesocosm: log10-ratio = −0.843 [−3.01, −0.222], *p* = 0.0094). This resulted in lower heritability for males than for females in both populations.

### Additive genetic variation in growth

The decomposition of the additive genetic variance of growth curve showed that the heritability of growth curve conditional to its average shape (see Methods, 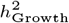) was small to moderate for both populations (Table 5). The difference in heritability between males and females was larger in the wild population, compared to the mesocosm population (Table 5). The dynamics of heritability along age was different depending on sex and population (Figure 3), although the uncertainty in the estimation was too large to attain significance in the wild population (maximal log10-ratio = −0.636 [−1.22, 0.0629], *p* = 0.0580, at age 1), but heritabilities in females were significantly higher for the mesocosm population after age 3 (log10-ratio = 0.754 [0.0721, 1.81], *p* = 0.0280 at age 3).

**Table 5.** Estimates obtained from the additive genetic variance decomposition of the growth curves using the Reacnorm package, depending on the sex (Females v. Males) and ages to average over (all ages, or adults of age ≥ 2 only). The total additive genetic variance of growth *V*_Add_ is decomposed into the additive genetic variance of intrinsic size *V*_A_ (genetic variation in being smaller or larger than other individuals on average across ages) and the additive-by-age variance *V*_AxAge_ (genetic variance in the shape of growth curve). Dividing by *V*_Tot_ (total variance, without the variation due to the average, shape of the growth curve) provides the heritabilities of growth, intrinsic size and shape of growth curve (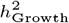,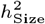 and 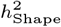). The *γ*- and *ι*-decomposition detail the relative contributions of genetic variation in each parameter to *V*_Add_ (total additive genetic variance in growth) and *V*_AxAge_ (additive genetic variance the shape of growth) respectively.

| Estim. | Wild |  | Mesocosm |  |
| --- | --- | --- | --- | --- |
|  | Females | Males | Females | Males |
| **Variances** |  |  |  |  |
| $V_{\text{Add}}$ | 1.39<br>[0.307, 2.63] | 2.62<br>[0.911, 4.53] | 3.11<br>[1.81, 4.63] | 2.48<br>[1.24, 4.48] |
| $V_A$ | 0.524<br>[0.218, 0.876] | 0.946<br>[0.396, 1.55] | 1.12<br>[0.766, 1.53] | 0.898<br>[0.599, 1.43] |
| $V_{\text{AxAge}}$ | 0.864<br>[0.0928, 1.77] | 1.68<br>[0.504, 3.02] | 1.99<br>[1.02, 3.07] | 1.57<br>[0.644, 3.06] |
| $V_{\text{Tot}}$ | 14.7<br>[12.1, 19.3] | 13.7<br>[11.0, 18.0] | 13.7<br>[10.5, 23.7] | 12.6<br>[9.14, 23.5] |
| **Heritabilities** |  |  |  |  |
| $h^2_{\text{Growth}}$ | 0.0941<br>[0.0202, 0.187] | 0.191<br>[0.0669, 0.334] | 0.225<br>[0.109, 0.336] | 0.192<br>[0.0905, 0.324] |
| $h^2_{\text{Size}}$ | 0.0353<br>[0.0138, 0.0639] | 0.0686<br>[0.0290, 0.114] | 0.0819<br>[0.0453, 0.117] | 0.0710<br>[0.0380, 0.106] |
| $h^2_{\text{Shape}}$ | 0.0586<br>[0.00583, 0.123] | 0.122<br>[0.0378, 0.222] | 0.143<br>[0.0626, 0.222] | 0.121<br>[0.0500, 0.220] |
| **γ-decomposition** |  |  |  |  |
| $\gamma_{L_0}$ | 0.151<br>[0.0676, 0.652] | 0.0792<br>[0.0398, 0.211] | 0.175<br>[0.113, 0.292] | 0.222<br>[0.123, 0.433] |
| $\gamma_k$ | 0.146<br>[3.75E-4, 0.737] | 0.767<br>[0.280, 0.934] | 0.343<br>[0.0884, 0.618] | 0.695<br>[0.329, 0.850] |
| $\gamma_{L_\infty}$ | 0.672<br>[0.0213, 0.888] | 0.150<br>[7.65E-4, 0.600] | 0.478<br>[0.195, 0.712] | 0.0840<br>[5.48E-4, 0.316] |
| **ι-decomposition** |  |  |  |  |
| $\iota_{L_0}$ | 0.0157<br>[0.00665, 0.149] | 0.00780<br>[0.00360, 0.0243] | 0.0413<br>[0.0233, 0.0806] | 0.0520<br>[0.0230, 0.134] |
| $\iota_k$ | 0.167<br>[4.18E-4, 0.922] | 0.821<br>[0.300, 0.990] | 0.424<br>[0.118, 0.730] | 0.857<br>[0.491, 0.964] |
| $\iota_{L_\infty}$ | 0.810<br>[0.0349, 0.985] | 0.170<br>[8.75E-4, 0.686] | 0.532<br>[0.227, 0.829] | 0.0923<br>[5.73E-4, 0.410] |

**Fig. 3.**
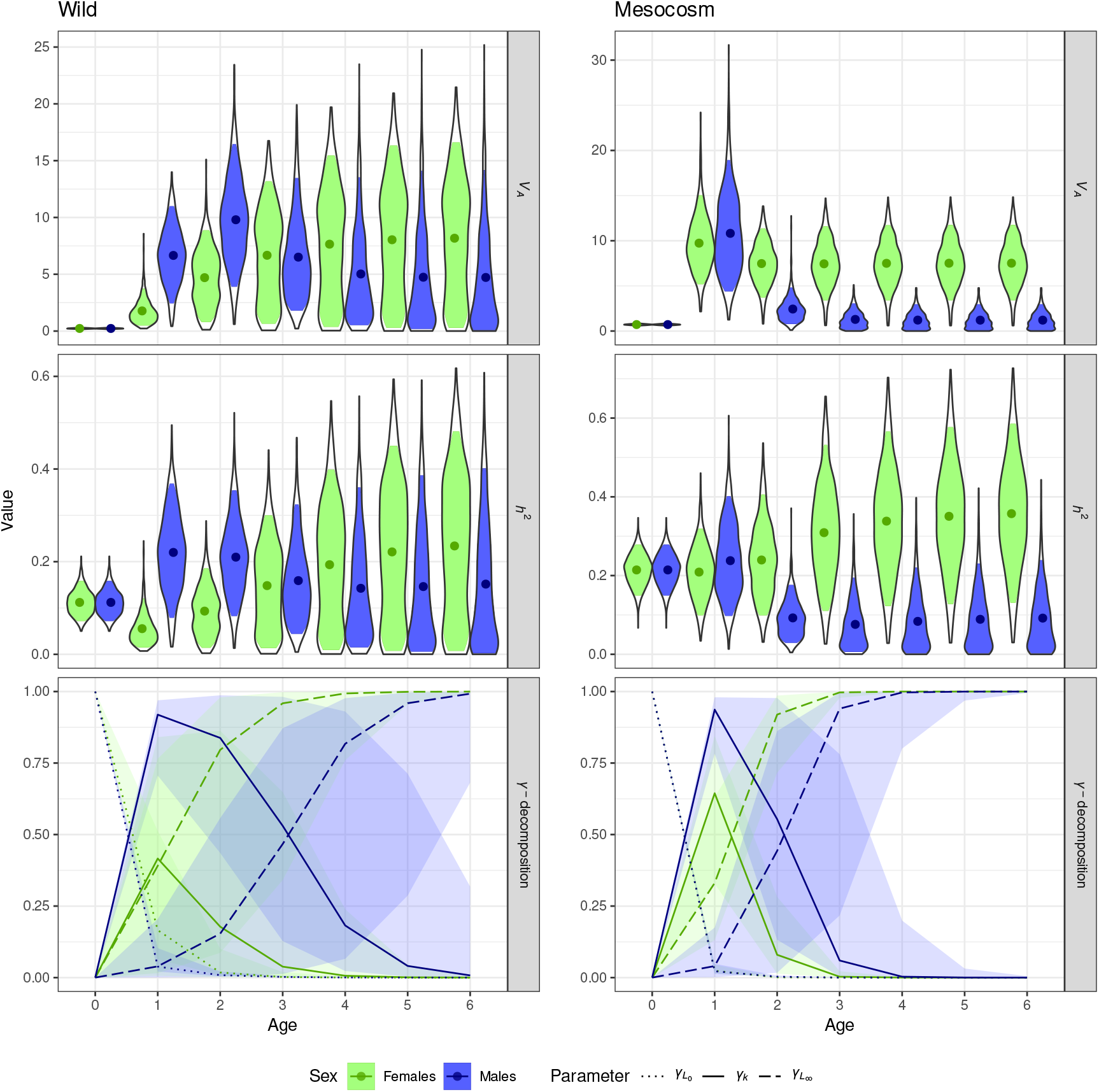
Additive genetic variance, heritability and *γ*-decomposition of the growth curve conditional to age and sex. The *γ*-decomposition shows, for each age, the contribution of the three parameters of the model (size at birth *L*_0_, growth coefficient *k* and asymptotic size *L*_*∞*_) to the additive genetic variance. Filling areas of the violin plots and transparent areas beyond the curves show the 95% credible intervals of the estimates.

The influence of the parameters in the additive genetic variation in growth (*γ*-decomposition of 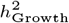, Table 5) strongly depended on sex and population: *k* was the most impacting parameter for males, while it was *L*_*∞*_ for the females. *L*_0_ played a minor to negligible role in the total genetic variation, although it was the second most important factor for males in the mesocosm population. The dynamics of the *γ*-decomposition against age reflected the biological interpretation of the parameters: while only *L*_0_ mattered at birth, *k* was the most important parameter in the earlier stages of life, then *L*_*∞*_ in the latter stages. The sex modulation of this dynamics reflected the sexual differences in the genetics of growth curves discussed above. Interestingly, the influence of the genetic variation in *L*_0_ was not entirely null at age 1 (Figure 3). Similarly, the influence of the genetic variation in *k* prolonged even after the asymptotic plateau was reached (age of 4 in the wild population and age of 2 in the mesocosm population). Comparing populations, the dynamics of the influence of *k* sharply reflected the observed difference in growth coefficients in Figure 2: its influence lasted for longer in the wild population than for the mesocosm population.

The heritability of intrinsic size 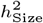 was very small (Table 5), though with statistical support away from 0, for all sexes and populations (but especially so for females in the wild population). The heritability of the shape of growth curve, 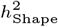 was larger than 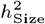 in all cases (Table 5), though not significantly so for the females in the wild population (log10-ratio = 0.153 [−0.397, 0.321], *p* = 0.276). Looking at the influence of each parameter to such genetic variation in the shape of growth (*ι*-decomposition, Table 5), the most important parameter was *k* for the males and *L*_*∞*_ for the females in both populations, following the pattern observed for the *γ*-decomposition. The importance of *L*_0_ for the additive genetic variance in the shape of growth was expectedly negligible.

## Discussion

To study the genetic bases of growth dynamics in a continuous growth species, we considered growth curve as a function-valued trait and analysed it with a non-linear animal model. We then applied a framework design to perform quantitative genetics on non-linear models (de Villemereuil and Chevin, 2025) to decompose the total heritability in growth curves (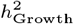) into the heritability of intrinsic size (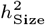, i.e. of being larger or smaller than other individuals on average) and the heritability of the shape of the growth curve (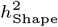). We were also able to disentangle the influence of the genetic variance in the parameters of the growth curve (i.e. initial size, asymptotic size and speed of growth) on such heritable variation (*γ*- and *ι*-decomposition).

### Environmental comparison of the populations

By comparing two populations of similar regional genetic origins in different environments, we were able to glimpse at the impact of the environment on lifetime growth dynamics. The environments varied between the populations in multiple ways (e.g. temperature, humidity, preys, predation) that a two-way comparison does not allow us to formally separate. We suggest nonetheless that thermal environment is a strong candidate to explain variation between populations. We found indeed, as expected from previous studies Chamaillé-Jammes et al. (2006); Bestion et al. (2015); Dupoué et al. (2017), that growth was faster in the warmer (mesocosm) environment, with a roughly doubled growth coefficient parameter. As a result, in the cold (wild) environment, individuals reach the asymptotic plateau only near the age of 4, meaning that very few individuals can reach such size before death. By contrast, in the mesocosm population, asymptotic plateau is reached sooner, at the age of 2.

Another difference between the populations was the shorter asymptotic size in the mesocosm population. When comparing different populations of the common lizard across a large-scale climatic gradient, Roitberg et al. (2013, 2020) found an overall decrease of female adult size with increasing winter temperatures. However, they found a positive slope between female adult size and winter temperature, when focusing specifically on the Western viviparous populations, the clade to which our populations belong. Similarly, using an experimental approach, Pellerin et al. (2022) found both an increase in juvenile growth rate in warmer environments, as we do here, and no impact of temperature on adult growth and adult size. Overall, the direct comparison between the results of these studies is made difficult by the heterogeneity of comparative/experimental approaches, and different characterisations of size, growth, climatic window studied and even definition of “adult” stage. Given the importance of size in the life-history of the common lizard (Sorci et al., 1996; Le Galliard et al., 2006, 2010; Bestion et al., 2015; Dupoué et al., 2017; Rutschmann et al., 2021), this environmental variation in growth curve is likely to impact most aspects of age-structured competition and selection in the population. In any case, these results are in agreement with both the metabolic theory of ecology (Brown et al., 2004; Huey and Kingsolver, 2019), predicting higher growth rate at higher ambient temperatures; and with the temperature-size rule (Atkinson, 1994; Atkinson and Sibly, 1997; Forster and Hirst, 2012), predicting that shorter asymptotic sizes are expected in warmer environments, with longer reproductive seasons (Adolph and Porter, 1996). In our case, the overall differences in growth curves between populations could be a direct effect of faster growth in the mesocosm population (*k* was twice as large than in the wild population), meaning that individuals have less time (i.e. fewer years) to accumulate resources to grow to bigger final sizes. This notably reflects the historical growth and development rates mismatch hypothesis to explain the temperature-size rule (van der Have and de Jong, 1996; Forster and Hirst, 2012). This alignment between theoretical expectation, previous results on the same species and the results exposed in this study favours the hypothesis that thermal environment is an important explanatory factor when comparing both populations. As mentioned above however, we cannot formally separate this factor from other possible environmental factors. Alternatively, the observed differences between populations could be explained by other life-history traits such as access to early reproduction (frequently at age 1 in the mesocosm population, Bestion et al., 2015) or impact of temperature on breeding phenology (Le Galliard et al., 2010; Rutschmann et al., 2016b; Massot et al., 2017) and reproductive investment (Rutschmann et al., 2016a); or any environmental differences between the populations other than temperature (though possibly correlated with it), such as humidity (Lorenzon et al., 2001), the quality of preys, or even genetic differences (Le Galliard et al., 2006).

### Quantitative genetics of growth

Using the residual variance for *L*_0_ and the permanent environment variance for *k* and *L*_*∞*_, as well as the additive genetic effect for all the parameters, we were able to distinguish between heritable and non-heritable components of individual variation, summarised in their heritability. We found heritable variation in all parameters of the growth curves, with the exception of growth coefficient *k* for females in the wild population, for which the estimate was very low. Heritabilities depended largely on a complex interaction between sexes and populations, especially for *k*. In all cases, as often is the case with heritability of traits in wild populations (Postma, 2014), most of the individual variation in the parameters was driven by environmental (or non-heritable genetic) components. We envision possible sources of such variation in the next sections. Interestingly, though expectedly when looking back at Figure 1, the influence of each parameter tends to spread beyond its direct influence on the growth curve: genetic variation in size at birth still has a small impact on individuals of age 1 and genetic variation in growth coefficient still impacts genetic variation in size after the asymptotic plateau is reached. Without a quantitative analysis like our *γ*-decomposition, it could thus be hard to circumscribe the ages affected by each parameters of the model.

Such heritable variation resulted in a moderate heritability of growth dynamics around 0.2, again with the exception of females in the wild population. Our framework allow us to decompose such heritability into the heritability of intrinsic size (“do individuals genetically vary in their tendency to be consistently larger or smaller than others across ages?”) and heritability of the shape of growth curve (“do individuals genetically vary in their growth rate, for a given general size?”). We found that, generally, two-third of the heritability of growth dynamics came from variation in their shape and only one-third in variation from their intrinsic size across ages. This means that genetic variation comes mostly from genetic variation in the growth strategy across individuals, rather than some individuals being larger or smaller across their lifetime.

### Environmental and genetic sources of sexual dimorphism

Size is an important ecological trait for both males and females in the common lizard, but partly for different reasons (Le Galliard et al., 2006; Horváthová et al., 2013a; Roitberg et al., 2013, 2020): while the advantage lays in competition for a territory or early access to reproduction in males (Gvozdík and Damme, 2003), it is more directly linked to fecundity in females (Horváthová et al., 2013b; Roitberg et al., 2013, 2020; Jurczyk and Borczyk, 2022). Especially, males to tend have lower adult survival and shorter lifespan than females in this species (Massot et al., 2011). This results in a sexual dimorphism in size and growth in this species, with males tending to be smaller than females (Le Galliard et al., 2006; Roitberg et al., 2020).

Our models confirmed a well-known pattern of sexual dimorphism in this species (Le Galliard et al., 2006; Roitberg et al., 2020). With our approach, we were able to examine this dimorphism over the lifetime of individuals and across two contrasted environments. Expectedly, the dimorphism rather lies in adult ages, and is thus more profoundly observed in the asymptotic size (*L*_*∞*_). The difference in the average asymptotic size was also found to be more intense in the mesocosm population, but there was a difference in the variance of asymptotic size between sexes, with males being less variable (Table 4). This result is consistent with the findings of Roitberg et al. (2020) that adult sizes of males are less variable across different populations.

This sexual dimorphism is also inscribed in the heritable genetic component. The sex-dependent additive genetic variances in the model allowed us to find a considerable difference between males and females in their genetic variation of growth, mostly reflecting the dimorphism described above, i.e. that males should be more constrained by competition at intermediate ages, while females should be more constrained by fecundity at later stages, strongly linked to their asymptotic size. On the one hand, we found that the heritable variation among males was mostly driven by their genetic variation in the growth coefficient (*γ*_*k*_ above 0.69), which was especially influential at earlier ages. On the other hand, we found that the heritable variation among females was mainly driven by the genetic variation in their asymptotic size (*γ*_*L*_*∞*__ above 0.47), which was especially influential at later ages. This has consequences, as it means that males and females are differentially responsive to selection on size at different stages on their lives, which could be the source of sexual conflicts. When comparing estimates across populations, the general levels of heritabilities, at all levels, were very comparable between populations for males (with a small exception for size at birth). By contrast, estimates for the females were different between the two populations: the estimate of the heritability of growth dynamics was smaller than males in the wild population, and larger than males in the mesocosm population. The most important difference between both populations for females was the very low estimated heritability for the growth coefficient found in the wild population, which strongly affected the dynamics of heritability across ages, especially when comparing heritabilites at age 1 between the two populations. Yet, in the wild population, most (56%) of the females reaching recruitment at the age of 2 breed only once. This means that the vast majority of the females never reach ages with the highest heritabilities, explaining the low overall heritability observed in this population. As mentioned above, males tend to die even earlier (77% of males reaching age 2 are not consequently observed), but their adaptive potential was also located in earlier stages of life (Table 4, Table 5, Figure 3). The reason for why the sexual contrast in the dynamics of heritability seems to behave differently between both populations can be linked to the *γ*-decomposition (Figure 3): the influence of *k* (i.e. *γ*_*k*_) is closer between sexes in the mesocosm population compared to the wild population, with a steeper decline with age in its relative influence against *γ*_*L*_*∞*__. Most of the difference between both populations in the dynamics of their heritability seem to stem from there and the values of 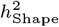. Two main, non-exclusive hypotheses could explain these patterns. The first would be that of differentiation at the genetic level. Although the two populations share the same regional geographic origins, there could still be differences due to local adaptation in this area. This is complicated by the fact that the mesocosm population is a hybrid stock of different source wild populations. The second hypothesis would be that strong genotype-by-environment interaction, generating different growth curves in different environments, even for the same genotype. This second hypothesis is almost guaranteed to hold, given the known and strong influence of e.g. temperature on growth dynamics of squamates as discussed above. What is unknown is whether the first hypothesis does hold, and the relative importance of both hypotheses. Such influence of temperature is, at the very least, compatible with the differences in the influence of *k* we observe between the wild and mesocosm populations, i.e. larger influence of heritable variation in *k*, even for the females, in the mesocosm population. Unfortunately, the current sampling of this study is not appropriate to settle and quantify the relative role of each hypothesis.

### Ecological and evolutionary consequences

Size and growth strategies are key ecological traits for squamates, as it influences many parts of their life-history, their metabolism or their thermoregulation (Barbault, 1988; Brown et al., 2004, 2022). Here, growth (or “size across ages”) was moderately heritable. On the one hand, this confirms the strong importance of the environment, and probably phenotypic plasticity, in the growth strategy of individuals. A confirmation that is further supported by our comparison between two environmentally different populations, hinting at genotype-by-environment, as discussed above. On the other hand, this result also highlights the adaptive potential both in size and growth. Since roughly two-third of the total heritability in growth curves stemed from genetic variation in the shape of growth (i.e. dynamics over ages), our result rather highlighted adaptive potential in growth strategies, i.e. the possibility for evolution to select strategies with e.g. different growth rates. By contrast, heritable variation in the intrinsic size of individuals, i.e. their consistent tendency in being small or large, accounted only for a third of the total heritability. This unveils that adaptive potential lies in heritable variation in the dynamics of growth, rather than size itself. In a context where global warming seems to impact a species like the common lizard mainly by accelerating its pace-of-life, especially early growth (Bestion et al., 2015; Dupoué et al., 2017, 2022), this is, in our opinion, best analysed through the lens of the temperature-size rule (van der Have and de Jong, 1996; Forster and Hirst, 2012), which predicts quicker growth earlier in life, but lower asymptotic sizes in a warmer environment. Projecting the temperature-size rule to future warmer climates indicates that phenotypic plasticity should mostly modify the shape of the growth curve, through an increase of the growth parameter *k* and decrease of asymptotic size *L*_*∞*_, with various consequences on the life-history of lizards, especially on the lifespan and long-term fecundity of females, both depending on their size at older ages (Bauwens and Verheyen, 1987; Hofmann and Henle, 2006; Horváthová et al., 2013a). Given this, it should be expected that selection pressures would arise to limit the decrease in *L*_*∞*_ (i.e. bigger females at older ages would still have higher lifetime reproductive success, despite the impact of the temperature-size rule on the plastic component of growth). In such case, this could result in a counter-gradient adaptive response (De Jong, 1988), i.e. an evolution toward decreasing the breeding value for the growth coefficient *k* and increasing the breeding value of *L*_*∞*_ to genetically counteract the effect of the temperature-size rule on the plastic component. This could even result in evolutionary rescue in some populations, although this remains highly speculative at the moment. Indeed, given the observed and predicted pace of phenotypic response to warming (Bestion et al., 2015; Dupoué et al., 2017; Bestion et al., 2023), the impact on population dynamics (Dupoué et al., 2022; Pellerin et al., 2022) and the moderate heritabilities inferred in this study, it seems that the likelihood of such scenario for most populations of common lizards at the Southern margin is thin. This is especially true as the most important sex for population dynamics, the females, were inferred to have a more limited adaptive potential for the evolution of growth in our wild population.

### Methodology

In this study, we applied a non-linear animal model framework, combined with a framework to decompose the additive genetic variance of non-linear function-valued traits designed to study reaction norms. Applying such framework allowed us to fit a parametric growth curve, combining information from all ages together, and yet, be able to relate additive genetic variation in the parameters back to the phenotypic traits of size and growth. This approach is close to the approach used by Kar et al. (2023) to study the quantitative genetics of growth on the delicate skink (*Lampropholis delicata*), with the notable difference that they relied on a quadratic curve, which is a linear approach from the parameter perspective (see de Villemereuil and Chevin, 2025). Such quadratic approach is possible in the early stages of development studied by Kar et al. (2023), but high order polynomials (with a large number of parameters) would be required to fit the lifetime growth curve of most squamates, which is not practical. By contrast, our non-linear approach only required three parameters to fit the growth curves. Another strong difference of our study is that we used a full pedigree of a wild population of common lizards to estimate the heritability of growth. Animal model based estimates of wild population heritabilities are scarce on squamates, and to our knowledge, this study is the first to provide such estimates for age-specific size and growth in a wild population of lizards. This is important as, for example, our estimates of heritability of size at birth is substantially smaller than previous estimates based on sib analysis (Le Galliard et al., 2006) and even on animal model with smaller sample size in the mesocosm population (Bestion et al., 2023). Differences with estimates not using the animal model can be explained by these methods being more naive and generally biased (Kruuk, 2004; Wilson et al., 2010; de Villemereuil et al., 2013). Differences with Bestion et al. (2023) most likely come from the increase in sample size or modelling choices like accounting for the cohort effect in this study.

## Conclusion

We provided a comprehensive quantitative genetic analysis of growth curves in the common lizard, demonstrating significant genetic variation in growth parameters across two populations in different environments. By applying a novel framework of additive genetic variance decomposition, we have gained valuable insights into the genetic basis of growth strategies in this species. These findings highlight, at least in this species: *(i)* the existence of adaptive potential for the evolution of the shape of growth curves; *(ii)* a very different ecological and evolutionary context for males and females; and *(iii)* a strong influence of the environmental context on both growth dynamics and its genetic determinants. Further studies regarding selection and the eco-evo dynamics of growth are needed to better understand the putative role of the evolution of growth to respond to selection, e.g. in the context of climate change.

## Data Availability

The data and code necessary to replicate this study are available at https://github.com/devillemereuil/CodeLizardGrowth.

## Author contributions statement

P.d.V. designed the study, with help from A.R and J.C. A.A. and P.d.V. performed the statistical analysis. A.A., P.d.V., A.R., M.R. and J.Co. interpreted the results. A.R., M.R., M.M., J.Cl. and P.d.V. collected data for the wild population. E.B., F.P., L.W., L.-M.S.J. L.D.G., E.D., J.C. and J.Co. collected data for the mesocosm population.

M.R. produced the pedigree for both populations. A.A. and P.d.V. wrote the manuscript, with help of A.R, M.R. and J.Co. and contributions from all co-authors.

## Acknowledgments

We are indebted to Don Miles, Matthieu Brevet, Andréaz Dupoué, Qiang Wu, David Rozen-Rechels, Sylvain Moulherat, Virginie Lepetz and all the other people, who are too numerous over the years to be listed here, who participated to the field work both in the wild and mesocosm population. We thank the *Parc National des Cévennes* for their welcoming support of the wild population survey. The wild population survey is part of the long-term Studies in Ecology and Evolution (SEE-Life) program of the CNRS. This work is part of a project that has received funding from the European Research Council (ERC) under the European Union’s Horizon Europe research and innovation program (EvoGenArch, grant agreement no. 101114976 to P.d.V.). P.d.V. was also supported by the *Institut Universitaire de France*. This work is part of a project that has received funding from the European Research Council (ERC) under the European Union’s Horizon 2020 research and innovation program (ECOFEED, grant agreement no. 817779 to J.Co.). J.Co., E.B., J.Cl. and M.R. were also supported by the French Laboratory of Excellence project ‘TULIP’ (grant nos. ANR-10-LABX-41 and ANR-11-IDEX-0002-02) and by an ‘Investissements d’avenir’ program from the Agence Nationale de la Recherche (grant no. ANR-11-INBS-0001AnaEE-Services). A.A. internship was funded by the *Muséum National d’Histoire Naturelle*. Views and opinions expressed are however those of the author(s) only and do not necessarily reflect those of the European Union or the European Research Council Executive Agency. Neither the European Union nor the granting authority can be held responsible for them.

